# Proximodistal organization of the CA2 hippocampal area

**DOI:** 10.1101/331025

**Authors:** I. Fernandez-Lamo, D. Gomez-Dominguez, A. Sanchez-Aguilera, E. Cid, M. Valero, L. Menendez de la Prida

## Abstract

The proximodistal axis is considered a major organizational principle of the hippocampus. Interfacing between the hippocampus and other brain systems, the CA2 region apparently breaks this rule. Apart from its specific role in social memory, CA2 has been involved in temporal and contextual memory but mechanisms remain elusive. Here, we used intracellular and extracellular recordings followed by neurochemical identification of single-cells to evaluate CA2 and surrounding areas in the rat. We found marked proximodistal trends of synaptic activity, as well as in subthreshold membrane potentials and phase-locked firing coupled to theta and gamma oscillations. Opposite proximodistal correlations between membrane potential fluctuations and theta sinks and sources at different layers revealed influences from up to three different generators. CA2 memory engrams established after a social memory task reflected these trends. We suggest that the structure and function of CA2 is segregated along the proximodistal hippocampal axis.

## Introduction

The CA2 hippocampal region has distinctive molecular, physiological and connectivity properties (Dudek et al. 2016). CA2 pyramidal cells respond vigorously to direct entorhinal inputs from layer II stellate cells (Sun et al. 2017; Leroy et al. 2017). In addition, they receive a direct mossy fiber connection from granule cells and contribute to a parallel trisynaptic circuit to deep CA1 sublayers (Kohara et al., 2014; Sun et al. 2017). Recurrently associated with CA3, CA2 pyramidal cells also project to superficial layers of the medial entorhinal cortex (Rowland et al. 2013). GABAergic innervation by local parvalbumin-expressing cells and specific classes of dendritic-targeting interneurons is particularly prominent in this region (Mercer et al. 2012a,b; Botcher et al. 2014), supporting a strong inhibitory control (Chevaleyre and Siegelbaum 2010; Piskorowski and Chevaleyre 2013). Notably, CA2 is specifically targeted by hypothalamic fibers releasing vasopressin and oxytocin during social interaction (Cui et al. 2013; Caldwell et al. 2008) and by suprammamillary glutamatergic cells with major role in wake-sleep regulation (Soussi et al. 2010; Pedersen et al. 2017).

Given recurrent connections with these different brain systems, CA2 can be considered a critical network gateway. Not surprisingly it is involved in a diversity of functions, including spatial and social memory. Place fields are more abundant but less precise in CA2 than in CA3 and CA1 (Oliva et al. 2016a). Some studies have revealed that CA2 ensemble firing changes prominently over time (Lu et al. 2015; Mankin et al. 2015; Lee et al. 2015; Alexander et al. 2016). In contrast, others have reported some cells firing in place during brief exploratory pauses and over sleep (Kay et al. 2016). This leads to the idea that CA2 is specialized in bridging contextual representations across time, supporting their contribution to episodic memory function (Wintzer et al. 2014; Mankin et al., 2015). When CA2 cells are specifically manipulated, defects emerge in contextual habituation to a neutral environment (Boehringer et al. 2017) but not for contextual fear memory or spatial learning (Hitti and Siegelbaum 2014). Instead, memory for a familiar conspecific is affected. Such a role in social memory may reflect not only specific features of CA2 (Leroy et al. 2017), but also downstream effects over deep CA1 pyramidal cells (Okuyama et al. 2016). Possibly, CA2 is instrumental in balancing microcircuit operation across different brain systems, but the mechanisms are ignored.

Recently, using extracellular recordings able to segregate unit firing, heterogeneous behavior of putative CA2 cells during hippocampal oscillations was reported (Oliva et al. 2016a,b; Kay et al. 2016). Moreover, in evaluating proximodistal changes of firing similarity between contexts, researchers found a significant inhomogeneity close to the CA3a/CA2 border (Lu et al., 2015). Unfortunately, without morphological confirmation it is difficult to interpret these results given the miscellaneous composition of this transitional area (Valero et al. 2015). Here, we used intracellular and extracellular in vivo recordings followed by neurochemical identification to target single cells around CA2 in rats. We found marked proximodistal trends of synaptic activity and theta/gamma oscillations in both subthreshold membrane potentials and phase-locked firing. Our data disclose an opposing entrainment by different theta generators and GABAergic microcircuits at the proximal and distal sectors. Moreover, we found these trends reflected in CA2 memory engrams. We propose that CA2 operates as a distributed network gateway to contribute their different function across brain systems.

## Results

### Characteristic inhomogeneities of local field potentials recorded around CA2

Local field potentials (LFP) were recorded with multi-site silicon probes around CA2 in 5 spontaneously active head-fixed rats. To target CA2 precisely, we learnt to identify characteristic evoked responses to stimulation of either the ipsilateral perforant pathway (PP) or the contralateral CA3 (Supp.Fig.1A; Methods). Theta oscillations and sharp-wave ripples were recorded during periods of running and immobility, respectively.

We noted attenuation of theta activity and characteristic sharp-wave ripple patterns at the CA2/CA1 border, as identified by the specific marker PCP4 (Fig.1A). Immunostaining against calbindin (CB) helped us to further delineate the limit of mossy fibers (MF) and the transition to CA1 (Fig.1A; arrowhead; Valero et al. 2015). Over penetrations, the probe also targeted CA3a,b and CA1 proximal (CA1p) regions. In CA3a and CA3b, gamma oscillations were typically present (Fig.1A, right). Similar LFP profiles were recorded under urethane in 30 rats (Fig.1B), in spite of spectral differences with drug-free condition.

**Figure 1:**
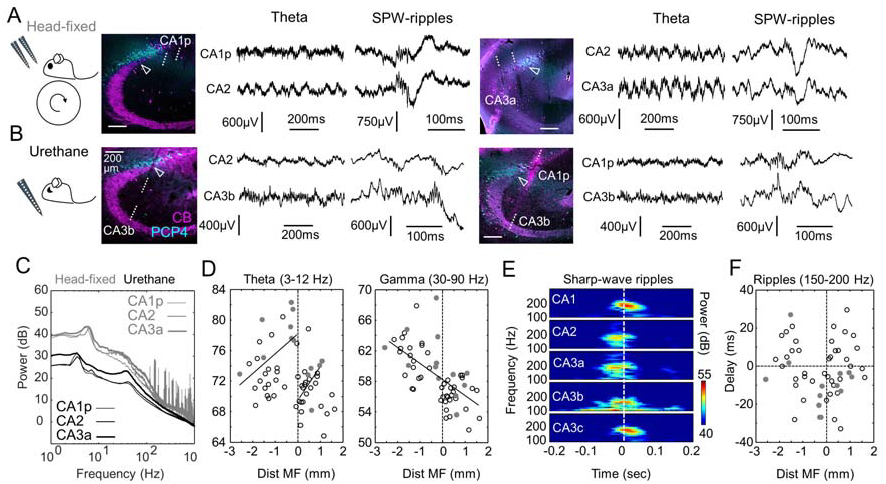
Characteristic in-homogeneities of local field potentials around CA2. **A**, Representative LFP signals recorded at SP in two head-fixed rats (left, right) using multi-site silicon probes. Probe tracks are in sections immunostained against PCP4 and CB. The limit of MF (open arrowheads) is taken as a reference for quantitative analysis. **B**, LFP recordings around CA2 obtained from urethane anesthetized rats. **C**, Representative power spectra during theta activity recorded at the SP of CA2, CA3a and CA1p under urethane (black) and in head-fixed conditions (gray). **D**, Individual spectral area of the theta (3-12 Hz) and gamma bands (30-90 Hz) plotted as a function of electrode distance to MF. Data from 52 recording locations, n=30 urethane anesthetized rats; n=12 recordings, n=5 drug-free rats. Different Pearson correlation was obtained at both sides of MF for theta: R=0.47, P=0.0059 from −3 mm to 0 and R=0.59, p=0.0088 from 0 to 1 mm. Gamma power exhibited significant negative correlation (R=-0.65, p<0.0001). **E**, Grand average spectra of the ripple power recorded at SP (aligned by the sharp-wave peak at SR). **F**, Delay between the ripple and SPW peaks as a function of recording location. Note earlier ripple peak at the CA3a/CA2 border.

We evaluated LFP features quantitatively using recording sites along the stratum pyramidale (SP) (Fig.1C). The spectral power of theta (3-12 Hz) and gamma bands (30-90 Hz) was plotted as a function of the site distance to MF (Fig.1D; n=12 recording sites from 5 head-fixed rats; n=52 recordings from 30 anesthetized rats; Methods). We noted representative LFP in-homogeneities around CA2. For theta, positive correlations appeared at both sides of the MF limit in an otherwise negative global trend (Fig.1D, left). This correlation paradox was not present in the gamma power, which decreased consistently (Fig.1D, right). We also confirmed characteristic features of sharp-wave ripples around CA2 when looking at the temporal relationship between the ripple power and the sharp-wave peak (Fig.1E). As recently described (Oliva et al. 2016b), we found the ripple power preceding the sharp-wave peaks at CA2 (Fig.1F).

Thus, distinctive features around CA2 suggest complex mechanisms might underlie the emergence of paradoxical LFP signals in a range of physiological events (theta, gamma and sharp-wave ripples).

### Cell-type specific heterogeneity around CA2

In order to investigate this question we characterized cellular diversity around this transitional hippocampal region. Immunoreactivity against PCP4, α-Actinin2, CB and Wfs1 allowed for classification of different cell-types (Methods). We found PCP4 immunoreactive cells at both sides of MF (Fig.2A, left), suggesting large dispersion of CA2 in rats as compared to mice (Fig.2B). Using the MF limit as a natural morphological landmark, we defined the proximal and distal sectors of CA2 (Fig.2A). We noted some PCP4+ cells in deep layers of CA3a (Fig.2A,B; open arrows). Similar to CA2, these cells co-expressed PCP4+ and α-Actinin2 (Supp.Fig.2). We also noted many Wfs1+ CA1 pyramidal cells interspersed with α-Actinin2+ cells at the distal CA2 region (Fig.2A, right). According to our estimations in rats, CA2 spans about 250 µm around the MF limit (anteroposterior −2.9 to −3.7 from Bregman).

**Figure 2:**
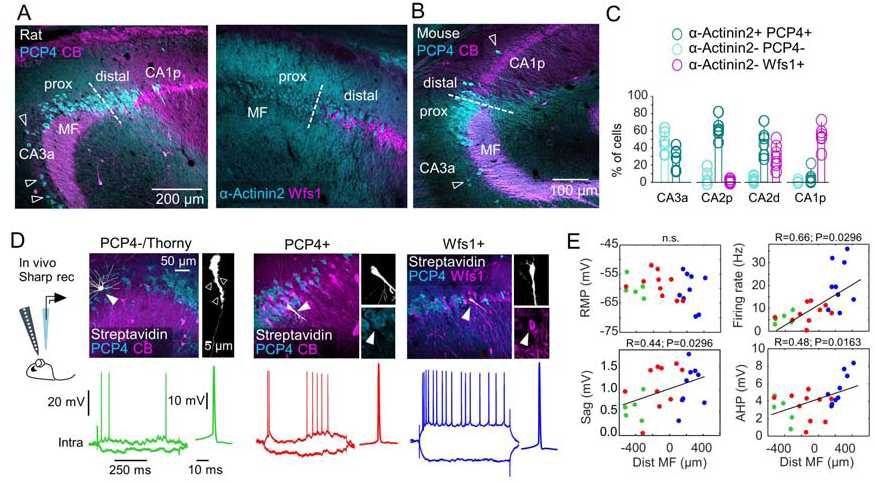
Cell-type specific heterogeneity around CA2. **A**, Immunoreactivity against PCP4, α-Actinin2, CB and Wfs1 allowed evaluating cell-type heterogeneity around CA2. Images show co-localization between different markers (3 confocal optical sections). Some PCP4+ cells were identified in deep layers of CA3a and in CA1 (open arrows). MF was used to define the proximal (close to CA3a) and distal (close to CA1) sectors of CA2. **B**, The dorsal CA2 of the mouse hippocampus exhibits a proximal organization. **C**, Quantification of distribution of pyramidal cells around CA2, as examined in double immunostaining against α-Actinin2/PCP4 (n=6 sections from 3 rats) and α-Actinin2/Wfs1 (n=6 sections from 3 rats). **D**, Intracellular recordings of CA2 pyramidal cells obtained in vivo from urethane anesthetized rats. Cells with thorny excrescences (open arrowheads) and lacking immunoreactivity for PCP4 or α-Actinin2 were classified as CA3 (green, n=5). Cells positive to Wfs1 either CB+ or CB-were classified as CA1 (blue, n=9). Cells immunoreactive to PCP4 or α-Actinin2 without thorny excrescences were classified as CA2 cells (red, n=10). **E**, Intrinsic properties of the different cell types plotted as a function of their distance to MF (n=24 cells from n=24 rats). Proximodistal gradients were confirmed by a Pearson correlation, as indicated. Intrinsic properties were measured at the resting membrane potential (RMP). Sag and firing rate were calculated in response to ±0.3 nA current pulses. AHP was calculated from the first spike in response to +0.2 nA.

Using VGAT-VenusA transgenic rats to exclude interneurons, we quantified the distribution of pyramidal cells with cell-type specific immunostaining. Double immunostaining against α-Actinin2/PCP4 (n=6 sections from 3 rats) and α-Actinin2/Wfs1 (n=6 sections from 3 rats) supported cellular heterogeneity (Fig.2C). At proximal CA2, most cells were PCP4+/α-Actinin2+ (∼65%) and some were negative for both markers (∼15%). These cells were VGAT-, indicating they had a glutamatergic phenotype. In double immunostaining against α-Actinin2/Wfs1 we found that a minority of these cells were Wfs1+ (<5%). In contrast, at distal CA2 many Wfs1+/α-Actinin2-cells (∼35%) intermingled with α-Actinin2+ cells (50%). The remaining Wfs1-/α-Actinin2-cells were VGAT+.

In order to identify different cell-types more precisely we performed intracellular recordings in vivo in urethane anesthetized rats in combination with 16-channel silicon probes (Fig.2D). We targeted pyramidal cells around the CA2 region using characteristic evoked responses to guide sharp electrode penetrations (Supp.Fig.1). After recordings, cells were filled with Neurobiotin, visualized with streptavidin and tested against PCP4, α-Actinin2, Wfs1 and CB immunoreactivity. Cell morphology was examined at 63x magnification under the confocal microscope to look for thorny excrescences, typical of CA3 pyramidal cells.

A total of 24 pyramidal cells were impaled in 24 rats. We found n=5 α-Actinin2-or PCP4-cells with thorny excrescences classified as CA3 cells (Fig.2D, green), n=10 cells expressing PCP4 being classified as CA2 pyramidal cells (red) and n=9 Wfs1+ CA1 pyramidal cells (blue). We recorded the proximodistal position of cell somata with respect to MF and confirmed the heterogeneous cellular composition of the CA2 region (Fig.2E). Electrophysiologically, we found no significant differences in the resting membrane potential (RMP) but proximodistal trends for the maximal firing rate and sag in response to current pulses, as confirmed by Pearson correlation (Fig.2E). We also noted some differences in the after-hyperpolarization (AHP) following an action potential (Fig.2E). Some of these features were early described in vitro for CA3a and CA2 pyramidal cells (Sun et al, 2017; Srinivas et al. 2017). While differences in input resistance between cell-types and difficulties to evaluate intrinsic properties in vivo may complicate interpretation (Supp.Table.1), altogether our data support heterogeneous cellular composition around CA2.

### Proximodistal differences of theta and gamma firing modulation around CA2

Intracellular recordings obtained simultaneously to multi-site LFP signals allowed us evaluating oscillatory behavior of identified cell-types (Fig.3A,B; n=24 cells). Juxtacellularly labelled cells recorded simultaneously to LFPs in head-fixed rats were also obtained (Fig.3B,C; n=3 cells; Methods). While theta under urethane (∼4Hz) may differ from running theta (6-7 Hz), we aimed to compare modulatory influences across cell-types.

**Figure 3:**
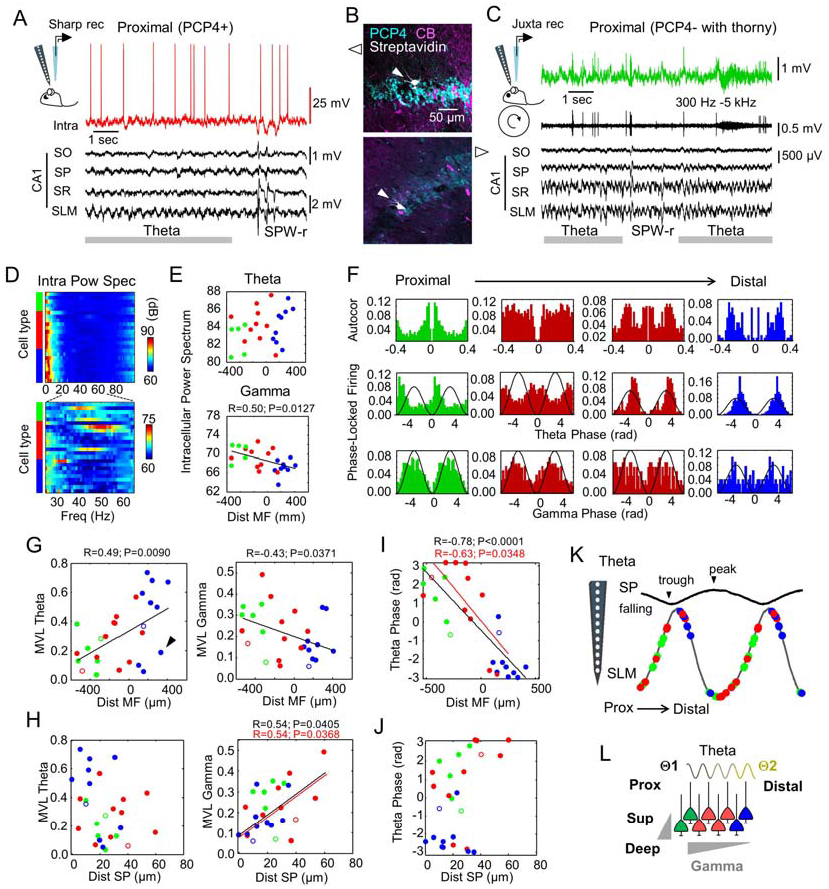
Proximodistal differences of theta and gamma activity of CA2 pyramidal cells. **A**, Intracellular recordings obtained simultaneously to multi-site LFP signals allowed evaluating oscillatory behavior of different cell types around CA2. Note poor theta rhythmicity of spontaneous firing of a prototypical PCP4+ CA2 cell, but consistent phase-locking preference with theta cycles at SLM. Note also clear hyperpolarization during SPW-ripples. **B**, Neurochemical classification of cells shown in A and C. **C**, Single-cell and LFP recordings from head-restrained rats. **D**, Power spectrum of the intracellular membrane potential recorded during theta in different cell types. Cells are ranked according to their proximodistal location within each group. Data from n=5 CA3 cells (green), n=10 CA2 cells (red) and n=9 CA1 cells (blue). **E**, Individual data of theta and gamma power of membrane potential oscillations. **F**, Representative examples of single-cell autocorrelation and phase-locking firing to theta and gamma waves recorded at the SLM. Cells are ranked according to their proximodistal location. **G**, Proximodistal distribution of the modulatory strength for theta and gamma for cells recorded under urethane (filled circles; 24 cells) and in drug-free conditions (open circles; 3 cells). Note separate cluster of poorly modulated cells (arrowhead). **H,** Distribution of the modulatory strength as a function of the cell distance within SP (0 is the superficial limit). **I**, Theta phase firing preference of single-cells measured against the SLM signal. **J**, Theta phase firing preference of cells plotted as a function of their deep-superficial location. **K**, Phase firing preference of single-cells represented against the CA1 SP signal (note reversal of theta wave along CA1 layers). **L**, Potential mechanisms may include proximodistal and deep-superficial microcircuit organization and influence of different theta generators.

Under urethane, the power spectrum of intracellular membrane potential oscillations suggested clear proximodistal gradients of the gamma power (30-90 Hz) across cells, but not clearly for theta (4-12 Hz) (Fig.3D,E; see Supp.Fig.3 for similar trends for slow and fast gamma band). Importantly, phase-locked firing of single-cells to LFP signals from the stratum lacunosum moleculare (SLM), exhibited opposite correlation for theta and gamma (Fig.3F). Theta-firing modulation increased towards distal CA2, and gamma modulation reversed, as quantified by the mean vector length (MVL; Fig.3G). For this analysis, we included juxtacellularly labelled cells, which fitted into the distribution (Fig.3G, open dots). We noted a separated cluster of poorly theta-modulated cells, especially some distal Wfs1+ and PCP4+ CA2 cells (Fig.3G, arrowheads), resembling in-homogeneities described before for LFP correlations.

Given deep-superficial gradients reported in CA1 and CA2 (Valero et al. 2015; Oliva et al. 2016b), we looked for the distribution of modulatory strength as a function of the cell distance within SP. We found significant correlation with the deep-superficial position for gamma modulation of all cells together and for the PCP4+ group alone (Fig.3H; superficial is at 0). A different modulatory index looking for pairwise phase consistency (PPC; Vinck et al., 2012), captured similar correlations including the CA1 pyramidal cell group (Supp.Fig.4A,B). Similar effects were seen for the slow and the fast gamma band separately (Supp.Fig.5). We noted however that due to the typical expansion of cell layer around CA2, the deep-superficial and proximodistal axes may interact. A generalized linear model (GLM) accounting for cell-types, proximodistal and deep-superficial influences, confirmed significant interactions (Supp.Table.2).

**Figure 4:**
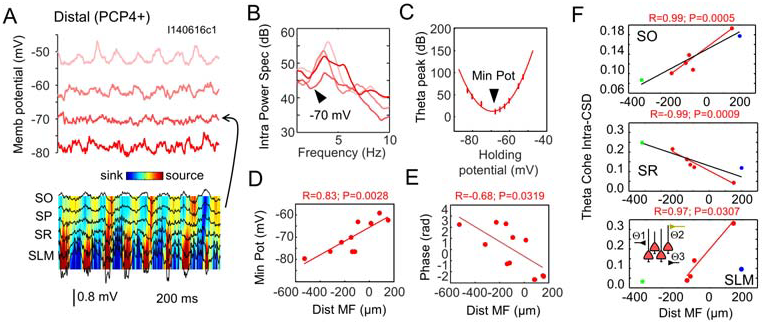
CA2 pyramidal cells couple to different theta generators along the proximodistal axis. **A**, Intracellular membrane oscillations recorded at different holding potentials simultaneously to extracellular LFP signals in one PCP4+ pyramidal cell. CSD local sinks and sources are shown together with LFPs (color map). Note attenuated theta oscillations at about −70 mV in this cell, near the reversal potential of GABAa receptors. LFP and CSD signals recorded simultaneously to the −70 mV trace are shown. **B**, Power spectrum of membrane potential oscillations of traces shown in panel A. Note reduced theta power for a holding potential near −70 mV. **C**, Relationship between theta power of membrane potential oscillations and the holding potential for the cell shown before. A minimum theta power is estimated at −70 mV (arrowhead). The thick line shows the best polymonial fit. **D**, Significant gradients of minimal power potential along the proximodistal axis. Data from n=10 PCP4+ CA2 cells. **E**, Phase relationship between membrane oscillation peak at RMP and the proximodistal location of CA2 cells. **F**, Proximodistal distribution of theta coherence between membrane potential oscillations at RMP and the local CSD signal at SO, SR and SLM. Data from cells recorded simultaneously to CA2 extracellular LFP signals (n=1 CA3, n=5 CA2, n=1 CA1). Inset shows schematically an intrahippocampal (SR) and entorhinal (SLM) theta generators (Θ1 and Θ2, respectively). A third independent generator likely contributes at SO (Θ3).

Importantly, along each theta cycle, proximal PCP4+ CA2 cells tended to fire in phase with CA3 cells, while distal CA2 cells clustered with Wfs1+ CA1 pyramidal cells (Fig.3I). Significant proximodistal gradients of the preferred theta phase were confirmed for all cells together and for PCP4+ cells alone (Fig.3I). The proximaldistal clusters emerged when plotted as a function of deep-superficial location, suggesting different deep-superficial dynamics (Fig. 3J). These effects did not depend on extracellular differences of LFP signals caused by penetrations of the silicon probe along CA2 to CA1 (Supp.Table.2; Supp.Fig.4C).

Altogether, our data suggest CA2 neuronal firing is segregated during theta oscillations. Considering the reversal profile of theta cycles around the SP of CA1, our data suggest that proximal PCP4+ pyramidal cells fire along the falling phase together with CA3 pyramidal cells, while distal cells fire in phase with CA1 cells at the theta trough (Fig.3K). Thus, firing from CA2 pyramidal cells consistently shifts in phase from proximal to distal. Complex mechanisms accounting for these functional effects could include local microcircuit factors and influence of different theta generators (Fig.3L).

### CA2 pyramidal cells couple to different proximodistal theta current generators

We sought to evaluate the contribution of different mechanisms with additional analysis. First, we focused in n=10 PCP4+ CA2 cells recorded intracellularly to avoid confounding factors due to cell-type heterogeneity. We examined intracellular membrane oscillations at different holding potentials during theta recorded extracellularly (Fig.4A). Current source density (CSD) signals allowed identifying theta-associated local transmembrane sinks and sources (Fig.4A, color map; Methods). Theta current generators were localized at different layers (Montgomery et al., 2009)

The frequency of intracellular theta oscillations was independent on the holding potential (Fig.4A), but the oscillatory power reached a minimum between −80 and −60 mV in different cells (Fig.4B, arrowhead). For a given cell, the intracellular theta power exhibited a characteristic U-shaped curve with a minimum theta power potential (Fig.4C). When plotted as a function of the cell position with respect to MF, the minimal power potential exhibited a significant proximodistal correlation for all PCP4+ cells (Fig.4D; no deep-superficial correlation, p=0.79). This minimal power potential reflected the equilibrium potential for mixed synaptic currents contributing to theta oscillations (Fig.4A; Soltesz and Deschenes 1993). At RMP, the phase of intracellular oscillatory peaks (Fig.4A, see trace at −50 mV) exhibited a significant proximodistal shift with respect to LFP theta (Fig.4F). These data suggest that CA2 pyramidal cells experienced different synaptic influences along the proximodistal axis during theta, with depolarizing peaks consistently shifting in phase from proximal to distal.

To examine this point further, we chose only those PCP4+ cells recorded simultaneously to CA2 extracellular LFPs (n=5). By evaluating coherence between the intracellular membrane potential and local CSD signals at stratum oriens (SO), radiatum (SR) and SLM, we aimed to quantify the influences of transmembrane currents at different strata during theta oscillations. We also included in this analysis one CA3 cell and one Wfs1+ CA1 pyramidal cell recorded simultaneously to LFP signals from CA2, to control for cell-type specific effects.

We found opposing proximodistal correlations between intracellular theta oscillations and CSD signals at SO and SR for all cells together and for PCP4+ cells alone (Fig.4E), suggesting that transmembrane currents flowing though basal and apical dendrites contributed differently along the proximodistal axis. Local CSD signals at SR likely reflect effects of intrahippocampal theta generators from CA3 (Montgomery et al. 2009) while SO CSD signals may reflect different contribution from extrahippocampal inputs (Buzsaki 2002). We also found positive proximodistal correlation for intracellular and local CSD signals at SLM but only for PCP4+ pyramidal cells (Fig.4E, bottom), consistent with their responsiveness to entorhinal inputs (Sun et al. 2017; Srinivas et al. 2017). Importantly, distal CA2 cells exhibited larger coherence with transmembrane theta currents at SLM than proximal CA2 cells.

Thus, our data suggest that different theta current generators influence proximal and distal CA2 cells. At proximal CA2, an intrahippocampal generator dominates while extrahippocampal inputs from the entorhinal cortex act distally across layers (Fig.4E, inset scheme). These different influences at the proximal and distal sectors of the transitional CA2 region may be responsible for paradoxical LFP correlations and functional gradients of theta and gamma modulation.

### Proximodistal gradients of synaptic responses along CA2

Next, we examine microcircuit determinants of proximodistal gradients within CA2 with stimulation of input pathways in vivo (Fig.5A) Intracellular synaptic responses were evaluated at different latencies in current-clamp for paired-pulse stimulation of the contralateral CA3 (n=20 cells) and ipsilateral entorhinal inputs (n=12 cells). Responses to electrical stimulation of entorhinal inputs were validated optogenetically (Supp.Fig.1).

**Figure 5:**
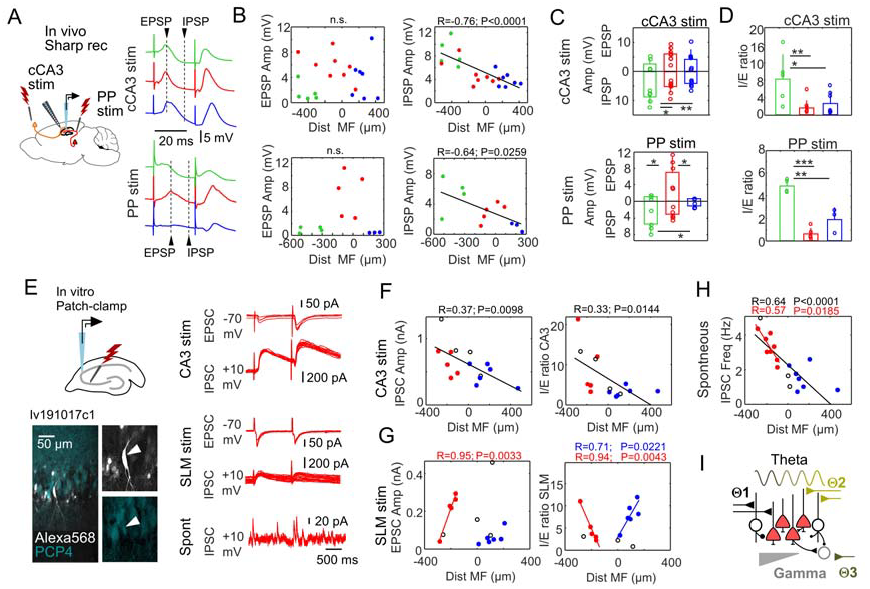
Proximodistal gradients of synaptic responses along CA2. **A**, Intracellular responses to cCA3 and PP stimulation were examined in vivo. The amplitude of evoked EPSPs and IPSPs were evaluated at different latencies from stimulation (arrowheads). **B**, Synaptic responses to cCA3 (n=20 cells) and PP stimulation (n=12 cells). Data plotted as a function of the cell distance to MF. **C**, Mean group responses per cell type. Note that due to their location, cell-type differences reflect a proximo-distal gradient along CA2. cCA3 stimulation: EPSP non-significant; IPSP F(19)=9.1, p=0.011, one-way ANOVA. *, p<0.05; **, p<0.005 post-hoc Tukey test. PP stimulation: EPSP F(11)=8.9, p=0.007; IPSP F(11)=6.1, p=0.021. *, p<0.05 post-hoc Tukey test. **D**, I/E ratio of different cell-types. cCA3 stimulation F(19)=6.5, p=0.008 one-way ANOVA. *, p<0.05, **, p<0.005 post-hoc Tukey test; PP simulation F(11)=41.1, p<0.0001. **, p<0.005, ***, p<0.0001 post-hoc Tukey test. **E**, In vitro recordings were obtained to evaluate synaptic currents in response to CA3 or SLM stimulation. Cells were filled with Alexa568 for posterior identification. Evoked excitatory (EPSC) and inhibitory (IPSC) synaptic currents from a PCP+ pyramidal cell are shown. **F**, Synaptic currents evoked by CA3 stimulation. Cells not confirmed neurochemically are indicated in black (n=4). Wfs1+ CA1cells (n=6) and PCP4+ CA2 cells (n=5) are shown in blue and red, respectively. Significant proximodistal trend for IPSC and the I/E ratio are indicated. **G**, Synaptic currents evoked by stimulation of entorhinal inputs at SLM (n=5 PCP4+ CA2 cells; n=6 Wfs1+ CA1 cells; n=4 not confirmed). **H**, Spontaneous IPSC frequency from n=18 cells (n=8 PCP4+; n=7 Wfs1+; n=3 not confirmed). **I**, Schematic representation of a proximodistal microcircuit organization of CA2

We found no proximodistal trends for the amplitude of excitatory potentials (EPSPs) elicited by stimulation of either pathway but significant correlation for di-synaptic IPSPs (Fig.5B). For entorhinal inputs, we noted clear EPSP responses only in PCP4+ cells, consistent with our results on theta coherence (Fig.4E, bottom) and in vitro data (Sun et al. 2017; Srinivas et al. 2017). Mean group responses per cell-type reflected these different trends (Fig.5C). An inhibitory/excitatory (I/E) ratio confirmed cell-type differences (Fig.5D). No effect was found for paired-pulse stimulation (Supp.Fig.6).

**Figure 6:**
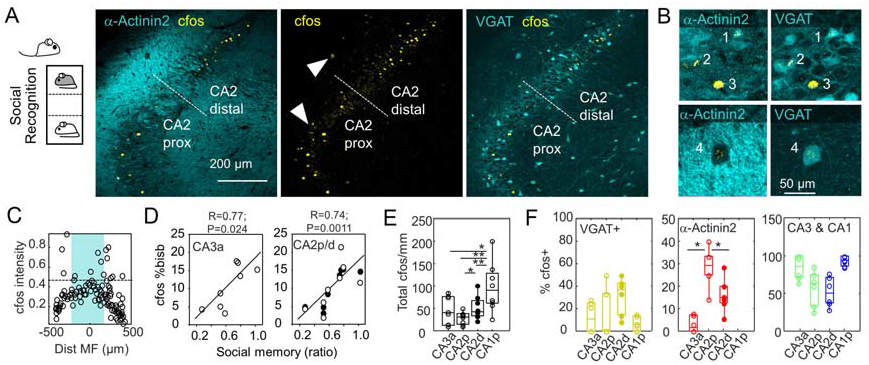
Proximodistal variability and heterogeneous composition of CA2 engrams. **A**, Rats were tested for their recognition memory of a familiar versus a novel conspecific. α-Actinin2 and VGAT signals are shown in the same false color to facilitate interpretation. Arrowheads indicate regions expanded in B. Note cfos expression in interneurons at SO. **B**, Enlarged view of regions indicated in A by arrowheads. Note cell-type specific heterogeneity of social memory engrams: α-Actinin2+/VGAT+ cell 1; α-Actinin2-/VGAT-cell 2; α-Actinin2+/VGAT-cell 3 and α-Actinin2-/VGAT+ SO interneuron 4. **C**, Distribution of single-cell cfos normalized intentisy as a function of the proximodistal position (one optical section). A threshold was defined for cfos counting (discontinuous line). The CA2 region was defined by α-Actinin2 immunoreactivity (shadowed area). **D**, Correlation between social recognition memory and the percentage of cells expressing cfos in one optical section per animal. **E**, Quantification of the linear density of cfos+ cells (one stack of 70 µm thickness per animal). Significant effects F(3)=9.6, p=0.0223 Kruskal-Wallis. *, p<0.05 and **, p<0.01 for a post-hoc Wilcoxon test. Data from n=8 rats. **F**, Percentage of cfos+ per cell type as identified in VGAT-VenusA rats (n=6). No statistical effects for VGAT+ interneurons. Significant effects for α-Actinin2+ cells: F(3)=19.2, p=0.0002 (Kruskal-Wallis). *, p<0.05 for post-hoc Wilcoxon test. Data from CA3 and CA1 was not tested statistically, as trends reflect regional distribution of each cell type.

To exclude cell-type specific effects and to examine synaptic currents more precisely we performed voltage-clamp patch-clamp recordings in vitro (Fig.5E, left; Methods). Cells in the vicinity of CA2 were recorded and filled with Alexa568 for neurochemical identification. Evoked excitatory (EPSC) and inhibitory (IPSC) synaptic currents were evaluated in response to CA3 and SLM stimulation (Fig.5E, right). We confirmed similar population responses for electrical and optogenetic stimulation of SLM (Supp.Fig.1D). In a preliminary set of experiments, we noted a significant effect of recording duration and access resistance in immunoreactivity against PCP4+, presumably due to cell dialysis (Supp.Fig.7). Thus, we tried to reduce patching time at minimum to gain in post hoc neurochemical characterization. Cells were first tested against PCP4 and subsequently for Wfs1. Cells without thorny excrescences and negative to both markers were left unclassified.

Consistent with in vivo data, we found significant proximodistal trends for the amplitude of IPSCs upon stimulation of CA3 (Fig.5F), but not for EPSCs with the exception of Wfs1+ CA1 pyramidal cells as tested in n=5 PCP4+, n=6 Wfs1+ and n=4 not-confirmed cells (Supp.Fig.8A). The I/E ratio also exhibited a significant correlation with the cell distance to MF (Fig.5F; similar trends for E/I: R=0.28, p=0.0237). Time to peak of evoked IPSCs was consistent with di-synaptic inhibition (Supp.Fig.9).

In contrast, responses to entorhinal inputs at SLM were more complex. First, we found significant proximodistal trends for the ESPC amplitude in PCP4+ cells (Fig.5G, left) and no differences for IPSCs (Supp.Fig.8B). Second, the I/E ratio showed opposing correlations for PCP4+ CA2 and Wfs1+ CA1 pyramidal cells (Fig.5G, right; similar for the E/I ratio; not shown). Interestingly, the frequency of spontaneous IPSCs, but not that of ESPCs, exhibited significant proximodistal correlation for all cells together and for PCP4+ cells (Fig.5H), with no difference in the amplitude of spontaneous events (Supp.Fig.8C). We found no clear evidence of deep-superficial gradients in evoked potentials in CA2 (Supp.Fig.10A). We also confirmed that MF stimulation elicited smaller EPSC responses in PCP4+ CA2 cell as compared with CA3a and larger I/E ratio (Supp.Fig.11), as previously reported (Sun et al., 2017).

Altogether, our data suggest that different theta generators converge along the proximal and distal sectors of CA2. The intrahipocampal generator running through CA3 pyramidal cells di-synaptically engages local GABAergic interneurons with stronger inhibitory influences at proximal CA2, consistent with gamma gradients. Instead, extrahippocampal theta generators have stronger influence at distal CA2 likely involving more complex mechanisms (Fig.5I). These different mechanisms may explain paradoxical LFP correlations in CA2 and different proximodistal modulation of CA2 pyramidal cells during theta activity.

### Proximodistal variability and heterogeneous composition of CA2 engrams

We reasoned that a proximodistal segregation of CA2 activity should be reflected in cognitive operations of this hippocampal region. To investigate this point, we tested rats for their recognition memory abilities of a familiar versus a novel conspecific, given a major role of CA2 in social recognition (Fig.6A; n=8 rats). Animals were perfused 1 hour after the task and coronal hippocampal sections were immunostained against cfos and α-Actinin2. GABAergic (VGAT+) cells were complementarily evaluated in a subset of VGAT-VenusA transgenic rats (Fig.6A,B; n=6). Expression of cfos was quantified separately in CA3a, proximal and distal CA2 and proximal CA1 (Fig.6C).

Rats submitted to the social recognition task exhibited different percentage of cfos+ cells in CA3a and CA2 as a function of their social memory ability (Fig.6D). For CA2, this association was significant for proximal and distal cells together (R=0.74; p=0.0011; open dots) and for the distal sector alone (R=0.85; p=0.0081; Fig.6D, filled dots). Non-significant correlation was found for cfos expression in CA1 (p=0.241). Quantification of the linear density of cfos+ cells confirmed significant regional effects (Fig.6E). Post-hoc paired tests showed more memory engrams at distal than proximal CA2 (Fig.6E).

To analyze the cellular composition of CA2 memory engrams, we separated the contribution from interneurons using sections from VGAT-VenusA rats and estimated the percentage of contributing cells from each type (Fig.6F; n=6 rats). Statistical effects were detected for α-Actinin2+ pyramidal cells, that were more engaged at proximal than distal CA2 engrams (Fig.6F). Many VGAT+ interneurons expressed cfos around CA2, including cells in the SO (Fig.6B; see cell 4). Interestingly, CA3a and CA1 pyramidal cells contributed to the engram at the proximal and distal sectors, respectively (Fig.6F, right). Thus, proximodistal segregation of activity is reflected in CA2 memory engrams supporting a cognitive role.

## Discussion

Our data suggest that within CA2, opposite influences from different theta current generators and local gradients of the excitatory/inhibitory ratio shape neuronal firing. The intrahippocampal theta converges proximally at SR to modulate CA2 pyramidal cell firing in phase with CA3a cells. In contrast at distal sectors, extrahippocampal activity operates maximally at SO and SLM to shift CA2 pyramidal cell firing towards CA1. The depth of theta modulation increases towards distal CA2, while gamma influences are stronger at proximal cells. These opposing trends interacted with differently assorted pyramidal cell-types, contributing to paradoxical LFP correlations. CA2 memory engrams established after a social memory task reflected such miscellaneous cellular composition and functional effects. Therefore, the CA2 output dissociates proximodistally in the dorsal hippocampus of the rat.

The transverse axis is central to hippocampal function. Initially identified in single CA3 pyramidal cells projecting to CA1 (Li et al., 1990; Isizhuka et al. 1990), a proximodistal topography quickly emerged as a major organizational principle of intra-hippocampal connectivity (Witter et al. 2000). Proximal CA3c cells (near DG) project distally to CA1, whereas distal CA3a project proximally. Analogous connectivity was identified for the CA1 projections to subiculum, where strong proximodistal gradients support functional dissociation (Amaral and Witter 1989; Cembrowski et al., 2018). Strikingly, the medial and lateral entorhinal inputs separate proximodistally in CA1 (Witter et al. 2000) and this is reflected in the organization of place fields (sharper at proximal CA1 and unspecific at distal; Henriksen et al., 2010). Similar functional segregation is present at CA3 (Lu et al. 2015). Given differences in recurrent connectivity (Ishizuka et al. 1990; Li et a., 1994), the ability of the CA3 network to separate patterns degrades towards CA3a, where more recurrent collaterals favor pattern completion (Lee et al. 2015).

Our data reveal that different proximodistal microcircuits determine functional properties of CA2 pyramidal cells as well. Distal CA2 cells receive stronger entorhinal inputs and lower di-synaptic inhibition in response to both CA3 and PP stimulation. We found that during spontaneous theta activity, the peak and reversal of membrane potential oscillations shifted in phase, consistent with different phase-locking preferences at proximal and distal sectors. Other report also noted different theta modulation in pyramidal cells recorded around CA2 in mouse, but neurochemical confirmation was not available (Matsumoto et al., 2016). Two independent analyses provided additional support to this functional segregation along the transverse axis. First, somatic membrane potential oscillations were more influenced by SLM and SO theta currents in distal PCP4+ CA2 cells, whereas proximal cells rather followed SR currents. Given the known layered organization of entorhinal, septal and supramammillary inputs at SLM and SO (Soussi et al., 2010; Joshi et al. 2017), our data suggest distal CA2 cells may be biased towards extra-hippocampal theta generators, while proximal CA2 cells rather follow intra-hippocampal CA3 influences. Proximodistal differences of I/E balance and unique plasticity properties of different pathways may further contribute (Piskorowski and Chevaleyre 2013; Nasrallah et al. 2016; Dasgupta et al., 2017; Leroy et al., 2017; Sun et al., 2017).

A second observation reinforced the idea of a proximodistal segregation of CA2 function. Gamma activity interfered largely with theta in proximal cells, as confirmed in both subthreshold membrane potential oscillations and neuronal firing. A confounding deep-superficial trend interacted with these proximodistal variations of feedforward inhibition, possibly reflecting diverse interneuronal connectivity (Mercer et al., 2012a,b; Botcher et al., 2014). Notably, axons from bistratified CA2 interneurons are shown to arborize distally while SP-SR interneurons innervate locally in proximal CA2 and CA3a (Mercer et al., 2012a,b). All these complex interactions along the proximodistal axis resulted in paradoxical LFP effects around CA2. While the gamma power measured at the SP consistently decreased, theta power exhibited a strong singularity at the point where MFs terminate. In addition to trends affecting CA2 pyramidal cells specifically, parallel extracellular dipoles from interspersed CA3a and CA1 pyramidal cells will contribute distinctly to LFP signals. Possibly, MF inputs recruiting preferentially CA3 versus CA2 cells (Sun et al., 2017; Kohara et al., 2014) and SLM inputs doing the opposite (Srinivas et al., 2017) will reinforce geometrical asymmetries in the area. Ripples preceding the local sharp-wave peak in the proximal, but not at distal locations, strengthen the idea of complex local LFPs explained by microcircuit mechanisms (Oliva et al. 2016b).

Blind recordings of CA2 cells have revealed contrasting activity during relevant physiological events. During sharp-wave ripples, some CA2 cells ramped slowly before exhibiting firing suppression while others discharge phasically prior to events (Oliva et al., 2016b). Using information from the vertical distribution of recording sites along silicon shanks, a deep-superficial organization was proposed for ramping and phasic CA2 cells, respectively. Tetrode recordings captured similar behavior during sharp-wave ripples (dubbed N and P cells, respectively) and extended some observations to theta (Kay et al., 2016). Interestingly, phase-locking distribution of P cells was slightly biased in both phase and modulation depth, as compared to N ‘ramping’ cells (see Extended Data Figure 6 of Kay et al., 2016). Putting these data in the context of our result, it is tentative to speculate that proximodistal effects of synaptic activity within the PCP4+ CA2 population, as well as contribution of interspersed CA3a and CA1 cells, can explain part of this functional diversity. Thus, in understanding CA2 function it will be critical to tease apart contribution of neighboring cell-types.

Spatial and non-spatial memories, as well as the ability for pattern separation, segregate along the transverse CA3-CA2 axis (Hunsaker et al., 2008; Nakamura et al., 2013; Lee et al., 2015). Place field rate differences between contexts drop abruptly at 200-250 µm from the CA2 border (Lu et al., 2015). This fits perfectly with the proximal CA2 region, where PCP4+ cells fire in phase with CA3a. At this border, oxytocin receptor signaling plays role in discriminating social stimuli (Raam et al., 2017). Consistently, we found significantly more α-Actinin2+ cells expressing cfos at proximal than distal CA2 after a social recognition task. We also noted a contribution from CA3a cells to social memory supporting the idea of critical CA3a/CA2 interactions (Ramm et al., 2017). Given recurrent interactions between CA3a-CA2 pyramidal cells (Li et al., 1994; Wittner and Miles, 2007), it is possible that a proximodistal integration of social and contextual information can be responsible for more flexible representations (DeVito et al., 2009; Wintzer et al., 2014; Pagani et al., 2015; Raam et al., 2017).

In contrast, distal CA2 cells fired in phase with proximal CA1 pyramidal neurons and a lower I/E balance suggest different computational operations as compared with the proximal sector (Guzman et al., 2016; Sun et al., 2017). Our finding that distal CA2 cells are more driven by entorhinal inputs than proximal cells suggest the circuit can accommodate additional functionalities (Jones and McHugh 2011). Consistently, we found Wfs1 pyramidal cells at the distal CA2 sector participating on memory engrams. Remarkably, social contexts can modify CA2 spatial fields (Alexander et al., 2016) possibly due to connectivity between CA2 and CA1 (Kohara et al., 2014; Okuyama et al., 2016). Importantly, we found differences between rats and mice regarding the extension of CA2 (Mercer et al., 2007; Kohara et al., 2014, San Antonio et al., 2014). Possibly, these differences reinforce the idea of microcircuit mechanisms supporting different social behavior across species (Chen and Hong 2018) and calls for caution on translational studies of psychiatric disorders.

In summary, we propose that CA2 operates along the proximodistal axis, similar to other hippocampal regions, and that this segregation is critical to better understand its functional role.

## STAR Methods

### KEY RESOURCES TABLE

See independent file

### CONTACT FOR REAGENT AND RESOURCE SHARING

Further information and requests for reagents and resources may be directed to the lead contact, Dr. Liset M de la Prida

### EXPERIMENTAL MODEL AND SUBJECT DETAILS

All protocols and procedures were performed according to the Spanish legislation (R.D. 1201/2005 and L.32/2007), the European Communities Council Directives of 1986 (86/609/EEC) and 2003 (2003/65/CE) for animal research, and were approved by the Ethics Committee of the Instituto Cajal (CSIC). Animals included in this study were not involved in any previous procedure.

A total of 58 adult (150 –200 g) and 16 juvenile (50-70 g) males and females rats were used (both wild-type and VGAT-VenusA transgenic Wistar rats). For in vivo electrophysiological experiments, 10 wild-type adult rats were used for head-fixed recordings and 30 for urethane anesthetized experiments. For in vitro studies, 14 wild-type and 2 VGAT-VenusA juvenile rats were used. For histological studies, 6 VGAT-VenusA transgenic rats were used. For optogenetic experiments, we used 4 wild-type adult rats injected with AAV-CamKII-ChR2. For behavioral assays, 2 wild-type and 6 VGAT-VenusA rats were used. All rats were maintained in the animal facility of the Instituto Cajal, with water and food *ad libitum* in a 12 h light-dark cycle (7am to 7pm).

### METHODS DETAILS

#### Juxtacellular and LFP recordings in head-fixed rats

Rats were first implanted with fixation bars under isoflurane anesthesia (1.5–2%) in oxygen (30%) while continuously monitored with an oximeter (MouseOx; Starr Life Sciences). After surgery, animals were habituated to head-fixed procedures (2-3 weeks habituation). The apparatus consisted in a cylindrical treadmill (40 cm diameter) adapted to a perforated table with a Narishige stereotaxic frame. The cylinder axle was equipped with a sensor to estimate speed and distance travelled analogically. The system was coupled to a water delivery pump controlled by a custom-made Arduino system. Animals learnt to run freely in the cylinder for water reward. After a couple of weeks of training, rats were able to stay confortable in the system for up to 2 hours with periods of running, immobility and sleep.

Once habituated to the apparatus, animals were anesthetized again to perform a craniotomy for electrophysiological recordings and stimulation (AP: −3.9 to -6 mm from Bregma; ML: 2-5 mm). A subcutaneous Ag/AgCl wire in the neck was implanted as reference and a bone screw served as ground. The craniotomy was sealed with sterile vaselyne and animals returned to their home cage. The day after, a bipolar tungsten wire was advanced to target CA3 while recording simultaneously from the contralateral CA1 with 16-ch silicon probes (NeuroNexus; 0.3–1.2 Mohm site impedance; 100 µm resolution; 177-413 µm^2^ electrode area) at the contralateral CA1. Once the stimulation position was adjusted, the electrode was cemented and the craniotomy sealed again with vaselyne. The day after (2 days after surgery for craniotomy) single-cell and LFP recordings started.

For LFP recordings of CA2, we used either a 32-ch silicon probe consisting in 2 shanks of 16-channels linear arrays each separated 200 µm or single 16-channels linear arrays (100 µm resolution; 413 µm^2^ electrode area). Penetrations were guided by characteristic response to cCA3 stimulation (Fig.1A; Supp.Fig.1). Then, single-cell recordings followed by juxtacellular labeling for post-hoc immunohistochemical identification were obtained in combination with LFP (Fig.3C). For juxtacellular recordings, a glass pipette (1.0mm × 0.58mm, ref 601000; A-M Systems) was filled with 1.5-2.5 % Neurobiotin in 0.5 M NaCl (impedance 8-15 MΩ). LFP signals were pre-amplified (4x gain) and recorded with a 32-channel AC amplifier (100x, Multichannel Systems) with analog filters (1Hz-5 kHz). Juxtacellular signals were acquired with an intracellular amplifier (Axoclamp 2B; Axon Instruments) at 100x gain. Single-cell and simultaneous LFP recordings were sampled at 20 kHz/channel with 12 bits precision (Digidata 1440; Molecular Devices).

After recording, cells were modulated using the juxtacellular labeling technique with positive current pulses (500-600 ms on-off pulses; 5-18 nA) while monitoring their response, as described before (Valero et al., 2015). After experiments, rats were perfused with 4 % paraformaldehyde and the brain cut in 70 µm coronal sections. Labeled cells were identified using streptavidin-conjugated fluorophores and submitted to immunostaining studies. Only unambiguously identified cells are reported (3 out of 10 cells).

#### Intracellular and LFP recordings under urethane

Procedures are similar to those described previously (Valero et al., 2015). Rats were anesthetized (urethane 1.2 g/kg, i.p.), fastened to the stereotaxic frame and kept warmed (37° body temperature). Bilateral craniotomies were performed for stimulation (cCA3 at AP: -1.2 mm, ML: 2.9 mm; ipsilateral PP at AP: −7 mm; ML: 3.5 mm), and a window was drilled above the right hippocampus for recordings (AP: −3.7 mm; ML: 3 mm). The dura was gently removed and the *cisterna magna* opened and drained.

LFP recordings were guided by extracellular stimulation and electrophysiological hallmarks to target either CA1 or CA2 region (Supp.Fig.1). LFP signals were acquired as described before. Concentric bipolar electrodes were advanced 3.5 mm with 30° in the coronal plane to stimulate CA3 or 3 mm vertically to stimulate PP. Stimulation consisted of biphasic square pulses (0.2 ms duration, 0.05-1.2 mA every 5 s). A subcutaneous Ag/AgCl wire in the neck served as reference. Recording and stimulus position was confirmed by post-hoc histological analysis. In most experiments, LFP were obtained from CA1 layers (n=4 CA3; n=5 CA2 and n=8 CA1). In an additional subset of experiments, single cells were recorded simultaneously to local LFP signals in CA2 (n=1 CA3; n=5 CA2 and n=1 CA1).

For intracellular recording and labelling in current-clamp mode, sharp pipettes (1.5 mm/0.86 mm outer/inner diameter borosilicate glass; A-M Systems, Inc) were filled with 1.5 M potassium acetate and 2% Neurobiotin (Vector Labs, Inc; impedances: 50-100 MΩ). Signals were acquired with an intracellular amplifier (Axoclamp 900A, 100x gain). Before recordings started, the craniotomy was covered by 3% agar to improve stability. The resting potential, input resistance and amplitude of action potentials was monitored all over experiments. After data collection, Neurobiotin was ejected using 500 ms depolarizing pulses at 0.5-2 nA at 1 Hz for 10-45 min. Rats were perfused with 4 % paraformaldehyde and the brain cut in 70 µm coronal sections for posterior histological studies.

#### In vitro electrophysiology

Juvenile wild-type and VGAT-VenusA transgenic Wistar rats were used to prepare hippocampal slices, when PCP4 was already expressed (San Antonio et al., 2014). Animals were anesthetized with pentobarbital, intracardially perfused with cold 95% O2 - 5% CO2 slicing artificial cerebrospinal fluid (ACSF) and decapitated using approved procedures. Sagittal slices (400 µm) were prepared from the dorsal level of the hippocampus at 2-4 mm from midline using a Leica vibratome (Leica VT1200S). Slices were cut and incubated for recovery at least 15 minutes at 32 ° C with a slicing ACSF whose composition was (in mM): 70 Sucrose, 86 NaCl, 2.5 KCl, 26 NaHCO3, 1

NaH2PO4, 0.5 CaCl2, 7 MgCl2, 25 glucose, pH 7.3 when balanced with 95% O2 - 5% CO2. After that, slices were transferred to a submerged holding chamber for at least 1hr at room temperature (RT) in recording ACSF containing (in mM): 125 NaCl, 2.5 KCl, 26 NaHCO3, 1.25 NaH2PO4, 2.5 CaCl2, 1.3 MgCl2, 10 glucose, pH 7.3 when balanced with 95% O2 - 5% CO2.

For recording, slices were transferred to a submerged recording chamber continuously perfused with ACSF (2.5-3 ml/min) using a peristaltic pump (Gilson) and oxygen-impermeable Tygon tubes. Somatic patch-clamp recordings were obtained from CA2 neurons under visual control with an upright microscope (BX51W, 60x lens; Olympus) at 32 ° C. Patch pipettes (pulled from borosilicate glass capillaries; World Precision Instruments, WPI) were filled with (in mM): 40 Cs-gluconate, 90 K-gluconate, 3 KCl, 1.5 NaCl, 1 MgCl2, 1 EGTA, 10 HEPES, 2 K2ATP, 0.3 NaGTP, 10 mM phosphocreatine and 0.1% Alexa568, pH 7.3 adjusted with KOH (osmolarity ∼300 mOsm). Electrodes filled with this solution had resistances of ∼4-6 MΩ.

Bipolar stimulating electrodes (tungsten wires, 0.5 mm separation, 0.5 MΩ, WPI) were positioned under visual control at the SP layer of CA3c/b or at the SLM layer of CA2. Extracellular field potentials were recorded with a patch-clamp pipette filled with ACSF coupled to one-channel AC amplifier (DAM-80; WPI). Stimulation intensity was adjusted homogenously between slices so that extracellular field potentials recorded at the CA1 or CA3 SR exhibited comparable responses (field EPSPs of 50-200 µV for 390-530 µA). IN a subset of experiments, MF were stimulated at the tip of the upper DG blade, known to project to the CA3a/CA2 border specifically.

Whole-cell patch recordings were obtained in current-and voltage-clamp modes with an Axoclamp 2B and digitized at 20 kHz (Digidata 1440A; Molecular Devices). Pyramidal cells had stable resting potentials of at least −50 mV and access resistances lower than 25 MΩ. Capacitance compensation and bridge balance were performed for current-clamp recordings. The junction potential was not corrected. After experiments, slices were fixed in 4% paraformaldehyde for histological studies.

#### Social recognition memory tasks

We used a social recognition memory task (Hitti and Siegelbaum 2014) to evaluate the proximodistal contribution of memory engrams around CA2. The apparatus consisted of a three-chamber box (central chamber: 20×50 cm; lateral chambers: 40×50 cm) situated in an evenly illuminated room with several cues visible on the surrounding walls. Male rats (150-180 g) were housed individually and had free access to food and water in a 12h light-dark cycle (7am to 7pm) at room temperature (21°C). Before the task, animals were habituated over 3-4 days to the empty apparatus and procedure (opening of guillotine doors; transferring to the inter-trial side box). Ambient noise was masked. Odor cues were removed after each trial (0.1 % acetic acid).

During the task, rats were placed in the center of the three-chamber box and doors opened. The task consisted of three phases of exploration, sociability and social novelty trials (10 min each). Each trial was followed by an inter-trial period of 10 min in which animals stayed in the familiar side box. During the exploration trial, two objects were located at the opposite chambers. During the sociability trial, a similarly aged male rat was placed in one chamber (randomized from rat to rat). The time spent exploring this rat versus the object was taken as an indication of sociability. During the social novelty test, a novel rat was placed in the other compartment and the time spent investigating each animal was scored.

Animal behavior was monitored with a video camera and analyzed offline. To quantify performance, we measured the time rats took to investigate objects and conspecifics, defined as an active approximation to each chamber (<2 cm). A recognition index was estimated for every object/rat by dividing the time difference taken exploring each element by the total exploratory time and tested against chance level (0.5; one-sample t-test). A recognition index of 1 indicates maximal interest for the conspecific versus the object (sociability index) or the novel versus de familiar conspecific (social memory). All rats exhibited normal sociability indices (object versus rat significantly >0.5). Animals exhibited different degree of social novelty (0.2-1). One hour after the task, rats were perfused with 4 % paraformaldehyde and brains cut in 70 µm coronal sections for cfos immunostaining.

#### Tissue processing and inmunohistochemistry

After completing experiments, animals were perfused with 4% paraformaldehyde and 15% saturated picric acid in 0.1 M, pH 7.4 phosphate buffered saline (PBS). Brains were postfixed overnight, washed in PBS and serially cut in 70 µm coronal sections (Leica VT 1000S vibratome). Sections containing the stimulus and probe tracks were identified with a stereomicroscope (S8APO, Leica). Sections containing Neurobiotin-labeled cells were localized by incubation in 1:400 Alexa Fluor488-conjugated streptavidin (Jackson ImmunoResearch 016-540-084) with 0.5% Triton X-100 in PBS (PBS-Tx) for 2 hours at RT. Slices recorded in vitro containing Alexa568 filled cells were fixed for 30 min, washed in PBS and processed similarly to others.

Sections containing the somata of recorded cells were treated with Triton 0.5% and 10% fetal bovine serum (FBS) in PBS. After washing, they were incubated overnight at RT with the primary antibody solution containing rabbit anti-calbindin (1:1000, CB D-28k, Swant CB-38), or mouse anti-calbindin (1:1000, CB D-28k, Swant 300) with 1% FBS in PBS-Tx to identify the MF. CB immunostaining was complemented with Wfs1 to identify CA1 pyramidal cells (1:1000, Proteintech 11558). For identifying the CA2 region, we used either rabbit anti-PCP4 (1:100, Sigma HPA005792) or mouse anti-α-Actinin2 (1:500;1:1000; Sigma A7811). CA3 pyramidal cells negative to Wfs1 and PCP4 were further examined for thorny excrescences. For cfos immunostaininig we used a polyclonal antibody at 1:100 (Santa Cruz Biotechnology sC-52. After three washes in PBS-Tx, sections were incubated for 2 hours at RT with secondary antibodies: goat anti-rabbit Alexa Fluor633 (1:500, Invitrogen, A21070), and goat anti-mouse Alexa Fluor488 (Jackson Immunoresearch 115-545-003) or goat anti-mouse Rhodamine Red (1:200, Jackson ImmunoResearch, 115-295-003) in PBS-Tx-1%FBS. Following 10 min incubation with bisbenzimide H33258 (1:10000 in PBS, Sigma, B2883) for nuclei labelling, sections were washed and mounted on glass slides in Mowiol (17% polyvinyl alcohol 4-88, 33% glycerin and 2% thimerosal in PBS).

Multichannel fluorescence stacks were acquired with a confocal microscope (Leica SP5; LAS AF software v2.6.0) and the following channels (fluorophore, laser and excitation wavelength, emission spectral filter) were used: a) bisbenzimide, Diode 405 nm, 415–485 nm; b) Alexa Fluor488, Argon 488 nm, 499–535 nm; c) Rhodamine Red / Alexa Fluor568 / Texas Red, DPSS 561nm, 571–620 nm; d) Alexa Fluor633, HeNe 633 nm, 652–738 nm; and objectives HC PL APO CS 10.0×0.40 DRY UV, HCX PL APO lambda blue 20.0×0.70 IMM UV, HCX PL APO CS 40.0x1.25 OIL UV and HCX PL APO 63x IMM OIL. For illustration purposes, false colors were used.

All morphological analyses were performed blindly to electrophysiological data. The lineal distance from the cell soma to the MF limit (taken as 0) or the cell position within SP (the superficial border taken at 0) was measured from confocal images using information from CB and bisbenzimide staining and the ImageJ software (NIH Image).

#### Optogenetics

To validate CA2 responses to electrical PP stimulation, we injected 2 wild-type Wistar rats with an adeno-associated virus (AAV5) carrying ChR2 under the control of CaMKII (AAV5-CamKII-ChR2) in the medial entorhinal cortex (Karl Deisseroth, UNC Vector Core). For stereotaxic surgery, rats were anesthetized with isoflurane (1.5–2%) in oxygen (30%). Injections of 1µl at AP −9 mm and ML 5 mm (DV 4 mm) were made unilaterally to the site of recording (titer 4.6 10^12^ vg/ml). Rats recovered for 2-4 weeks to allow for adequately expression. For in vivo optogenetic experiments (n=2 rats), light was delivered from a solid state blue laser (MBL-F-473, 300 mW maximal fiber output, CNI Laser, China) with Neuronexus opto-probes (flat fiber optic 105 µm diameter mounted on 16-ch linear arrays) located stereoataxically at the angular bundle. LFP signals were recorded simultaneously from CA2 and CA1 layers with an independent probe. Laser stimulation ranged from 3-15 mW for short pulses of 2-5 ms. After experiments, animals were perfused with 4% paraformaldehyde and sagittal sections (70 µm) were obtained to validate infection. In vivo experiments (n=2 rats), were performed similarly using an edged optical fiber (Optogenix; Italy) and 16-channel silicon combs from neuronexus (Prida design; interspaced 100 µm).

#### Analysis of LFP signals

Analysis of electrophysiological in vivo data was performed using routines written in MATLAB 7.10 (MathWorks). Multi-site LFPs from different layers were identified using characteristic physiological events, including sharp-wave ripples (to identify SR and SP) and maximal theta oscillations (for SLM). Characteristic evoked responses to contralateral CA3 and ipsilateral PP simulation were used to identify CA2 penetrations (Supp.Fig.1). One-dimensional current-source density (CSD) signals were calculated from the second spatial derivative of laminar LFPs (100 µm resolution). Smoothing was applied to CSD signals for visualization purposes only. Tissue conductivity was considered isotropic, and an arbitrary value of 1 was assigned to express CSD signal as mV/mm^2^.

The power spectrum of LFP signals was estimated using the Fast Fourier transform (FFT). For theta activity, non-overlapping segments of continuous oscillations in the 4-12 Hz band were identified in LFP signals from SLM. The contribution of theta (4-12 Hz) and gamma (30-90 Hz) activity was evaluated from the spectral area at each oscillatory band using data from recording sites at SO, SP, SR and SLM. Gamma activity was separated in the lower (30-60 Hz) and high (60-89Hz) bands. The location of probe penetrations along the SP was evaluated by estimating the linear distance to MF.

For sharp-wave ripples, LFP recordings from SR were low-pass filtered at 100 Hz to identify sharp-waves and signals from SP were bandpass filtered between 100-600 Hz to identify ripples. We used forward-backward-zero-phase finite impulse response (FIR) filters of order 512 to preserve temporal relationships. For sharp-waves, filtered signals were smoothed (Gaussian kernel) and events detected by thresholding of >3 SDs. For ripples, bandpass-filtered signals were smoothed (Savitzky-Golay) and events detected by thresholding of >2 SDs. All pairs of detected events were visually confirmed. Time-frequency analysis of sharp-wave ripples was performed by applying the multitaper spectral estimation in sliding windows with 97.7% overlap and frequency resolution of 10 Hz in the 90-600 Hz frequency range (only the 100-600 Hz range is shown) to data sweeps aligned by sharp-wave ripple events (± 1sec). The time difference between the sharp-wave peak and the maximal ripple power was estimated and plotted as a function of the probe position with respect to MF.

#### Analysis of intracellular recordings

Passive electrophysiological properties (input resistance, membrane decay and capacitance) of neurons recorded intracellularly in vivo were measured using 500 ms currents step in current-clamp mode. Cells with intracellular action potential amplitude smaller than 40 mV were excluded. RMP and input resistance were estimated by linear regression between baseline potential data and the associated holding current.

Intrinsic firing properties, including action potential threshold, half-width duration and AHP were estimated from the first spike in response to depolarizing current pulses of 0.2 nA amplitude and 500 ms duration. The sag and maximal firing rate was calculated from current pulses of ±0.3 nA amplitude. A bursting index was defined as the ratio of the number of complex spikes (minimum of 3 spikes <8ms inter-spike interval) over the total number of spikes recorded during theta activity.

The power spectrum of intracellular membrane potential oscillations was calculated with FFT methods similar to LFP signals for different holding potentials. The contribution of theta (4-12 Hz) and gamma (30-690 Hz) bands was evaluated from the spectral amplitude of the FFT. We found comparable results by using the spectral area below detrended power spectrum (not shown). Pairwise theta coherence between the intracellular membrane potential and LFP or CSD signals was defined from the cross-spectral power densities at the peak frequency in the 4–12 Hz range at 1Hz resolution.

Phase-locking firing of single cells was measured from each spike using the Hilbert phase of theta peaks recorded at SLM. Each theta cycle was divided into 25 bins. Phase locking was quantified using the mean vector length (MVL) of phase distribution from 0 to 1. We also used a pairwise phase consistency measure (PPC) suitable for evaluating modulation of relatively small number of spikes (Vinck et al., 2012). The SLM theta trough was set at 0 and peaks at ± pi. To establish links with previous data, we also estimated the corresponding theta peak at SP which exhibited a shift of -1.6 radians respect to the SLM trough due to theta wave asymmetries. To the purpose of this paper, this shift was not considered. Gamma modulation was evaluated similarly for LFP oscillations in the full gamma band (30-90 Hz) or in the slow (30-60Hz) and high (60-90 Hz) bands separately.

To evaluate theta rhythmicity of single-cell firing, we estimated the power spectrum of the autocorrelogram built at ±0.5 s windows (1 ms bin size). A theta autocorrelation index was defined from the normalized area in the 4–12 Hz band. For visualization purposes, 45 bins were used to build autocorrelograms.

#### Analysis of juxtacellular recordings

For juxtacellular recordings data from glass pipettes were high-pass filtered at 300 Hz to detect positive spikes from the juxtacellular recorded cell (> 8 SD). Simultaneous LFP signals at SLM and SP were processed similarly than for intracellular recordings. Interspike interval autocorrelograms (0.5 ms bins) were constructed using all detected spikes. The stability of the action potential waveform (peak-to-peak duration and amplitude as well as a spike asymmetry index defined as the ratio of the difference between the negative and positive baseline-to-peak amplitudes and their sum) was evaluated over the entire recording session (> 3 min), before juxtacellular electroporation. Baseline firing rate was stable for small movements of the pipette towards the cell, excluding mechanical interferences. Phase-locking firing was evaluated similar as described for intracellular data.

### QUANTIFICATION AND STATISTICAL ANALYSIS

Statistical analysis was performed with MATLAB. No statistical method was used to predetermine sample sizes, which were similar to those reported previously (Valero et al., 2015). Normality and homoscedasticity were confirmed with the Kolmogorov– Smirnov and Levene’s tests, respectively. The exact number of replications for each experiment is detailed in text and figures.

One-way ANOVAs or Kruskal-Wallis tests were applied for cell-types or regions. Post-hoc comparisons were evaluated with either the Tukey-Kramer or Wilcoxon tests. Proximodistal and deep-superficial trends were evaluated with the Pearson product-moment correlation coefficient, which was tested against 0 (i.e., no correlation was the null hypothesis) at p < 0.05 (two sided). Both the Pearson coefficient and p value are reported to facilitate interpretation.

To account for mixed statistical effects on measurements of interest, a generalized linear model (GLM) was implemented as a linear combination of the following variables: cell-type, distance to MF and distance within SP. The impact of each variable in the GLM model was then tested with ANOVA and the p value reported after Tukey-Kramer posthoc correction. Variables having a significant impact in explaining the measurement of interest show p values < 0.05.

### DATA AND SOFTWARE AVAILABILITY

Freely available software and algorithms used for analysis are listed in the resource table. Some analyses were specifically designed for the purpose of this paper using routines written in MATLAB 7.10 (MathWorks). All custom scripts and data contained in this manuscript are available upon request from the Lead Contact.

Author contribution
LMP designed the study. IFL, ASA, EC, MV obtained data.DGD, IFL, ASA, MV, EC, LMP analyzed and interpreted data. LMP wrote the paper.

## Acknowledgments

Supported by grants from the Spanish Ministerio de Economía y Competitividad (MINECO) to LMP (BFU2015-66887-R) and the Fundación Tatiana Perez de Guzman el Bueno. DGD and MV were supported by a PhD fellowship from the Spanish Ministry of Economy (BES-2013-064171) and from the Ministry of Education, Culture and Sports (FPU12/03776), respectively. We thank to the Deisseroth lab for sharing their optogenetic constructs. VGAT–Venus transgenic rats were generated by Drs. Y. Yanagawa, M. Hirabayashi, and Y. Kawaguchi at the National Institute for Physiological Sciences (Okazaki, Japan) using pCS2–Venus provided by Dr. A. Miyawaki. VGAT line progenitors were provided by the National Bioresource Project Rat (Kyoto, Japan)

